# Pervasive mRNA uridylation in fission yeast catalysed by both Cid1 and Cid16 terminal uridyltransferases

**DOI:** 10.1101/2022.10.04.510797

**Authors:** L Lipińska-Zubrycka, M Grochowski, J Bähler, M Małecki

## Abstract

Messenger RNA uridylation is pervasive and conserved among eukaryotes, but the consequences of this modification for mRNA fate are still under debate. Utilising a simple model organism to study uridylation may facilitate efforts to understand the cellular function of this process. Here we demonstrate that uridylation can be detected using simple bioinformatics approach. We utilise it to unravel widespread transcript uridylation in fission yeast and demonstrate the contribution of both Cid1 and Cid16, the only two annotated terminal uridyltransferases (TUT-ases) in this yeast.

To detect uridylation in transcriptome data, we used a RNA-sequencing (RNA-seq) library preparation protocol involving initial linker ligation to fragmented RNA. We next explored the data to detect uridylation marks. Our analysis shows that uridylation in yeast is pervasive, similarly to the ones in multicellular organisms. Importantly, our results confirm the role of the cytoplasmic uridyltransferase Cid1 as the primary uridylation catalyst. However, we also observed an auxiliary role of the second uridyltransferase, Cid16. Thus both fission yeast uridyltransferases are involved in mRNA uridylation. Intriguingly, we found no physiological phenotype of the single and double deletion mutants of *cid1* and *cid16* and only limited impact of uridylation on steady-state mRNA levels.

Our work establishes fission yeast as a potent model to study uridylation in a simple eukaryote, and we demonstrate that it is possible to detect uridylation marks in RNA-seq data without the need for specific methodologies.

## Introduction

Messenger RNA (mRNA) degradation has a critical role in regulating transcript levels and is a major point of gene expression regulation. Bulk cytoplasmic mRNA decay was most studied in budding yeast, where it is initiated by poly(A) tail shortening followed by removal of the 5’ cap (decapping) [1]. Removal of this protective cap structure renders mRNA accessible for the cytoplasmic 5’-3’ exonuclease Xrn1. In yeast, 5’-3’ decay is considered the main degradation pathway, however, mRNA can be also be degraded from the 3’ end by the exosome complex supported by the SKI complex [2]. While the degradation machinery is evolutionary conserved, progress in analysing other eukaryotic models demonstrated some peculiarities of the budding yeast system [3]. Most importantly, while pervasive cytoplasmic uridylation of mRNAs was detected in most studied eukaryotes it is absent in budding yeast [4,5].

Uridylation is catalysed by cytoplasmic terminal uridyltransferases (TUTases), while exonuclease Dis3l2 preferentially targets uridylated RNAs [6,7]. It is believed that transcript uridylation induces its degradation either by attracting the LSM complex which accelerates decapping and subsequent 5’-3’ decay or by triggering 3’-5’ decay by Dis3l2 exonuclease [4]. Important roles for this new regulation layer were identified in specific processes like antiviral response [8], transposon control [9], germline development [10,11], or RNA degradation triggered by apoptosis [12]. Still, the significance of pervasive mRNA uridylation in the context of bulk mRNA turnover is unclear, and elimination of uridylation generates no detectable consequences for the physiology of somatic cells and only limited molecular manifestations [7,11,13].

Messenger RNA uridylation was first detected in fission yeast *(Schizosaccharomyces pombe*) [5], another wellstudied single-cell eukaryotic model that is evolutionary distant from budding yeast. The fission yeast genome encodes two TUT-ase genes, *cid1* and *cid16* [14]. Cid1 was shown to be the main enzyme responsible for mRNA uridylation, while the contribution of Cid16 to this process is not known [5]. The *S. pombe* U-tail-specific exonuclease Dis3l2 was shown to genetically interact with mRNA decay factors suggesting its contribution to the bulk degradation process [7]. Uridylation in fission yeast was so far only detected on a few tested transcripts [5,7,15].

Currently, the only described method for genome-wide uridylation analysis is TAIL-seq. TAIL-seq involves specific library preparation that due to adapter ligation preserves 3’-end information; moreover, the TAIL-seq method involves a bioinformatics pipeline that improves the base-calling of the sequences including long adenine homopolymer [16]. TAIL-seq brought numerous new insights to our understanding of the eukaryotic mRNA degradation process, but it is a technically and computationally challenging method. RNA sequencing library preparation protocols involving 3’-adapter ligation to RNA after initial fragmentation also preserve some 3’-end information. Such information is usually used to map polyadenylation sites [17]. Here we used a fission yeast model to demonstrate that uridylation can be detected in such RNA sequencing data using a simple analysis pipeline. Our results show pervasive mRNA uridylation in fission yeast that shares characteristics with uridylation detected in higher eukaryotes. Moreover, we observe that both Cid1 and Cid16 uridyltransferases contribute to mRNA uridylation. Furthermore, we show that uridyltransferases deletion does not induce any detectable cell growth phenotypes and has only a limited impact on mRNA steady-state levels.

### Detecting 3’-end uridylation in RNA-seq reads

One way of RNA sequencing library preparation is through ligation of adapters to fragmented mRNA before library amplification. Sequencing data obtained from such libraries contain some reads with full or fragmented poly(A) tails, and such reads can be used to map the poly(A) sites of transcripts [17]. We reasoned that terminal uridylation could also be detected in this type of data (Fig. 1A). To check our idea, we prepared RNA sequencing libraries by ligating adapters to rRNA-depleted and fragmented total RNA using a previously described protocol [18]. Libraries were then subjected to Illumina sequencing, and the reads were aligned to the genome using an aligner with standard soft-clipping options (see Methods). Next, we used information encoded in the “.sam” file CIGAR string to identify reads with 3’-end non-templated nucleotides (Fig. 1A). In agreement with knowledge about mRNA uridylation, we looked specifically for reads with pure poly(A) or uridylated poly(A) extensions with uridines added at the 3-end of the poly(A) tail (poly(AU) reads). Additionally, we recorded reads with only U extensions or somewhat mixed A and U order. To minimise potential artefacts due to sequencing errors, we focused on the reads with extensions being at least 4 nucleotides long. The fraction of identified reads with 3’-end adenine and/or uridine un-templated extensions longer than 3nt accounted for 0.96% of total aligned reads (around 15 thousand reads in total for wild type sample).

**Figure 1.**
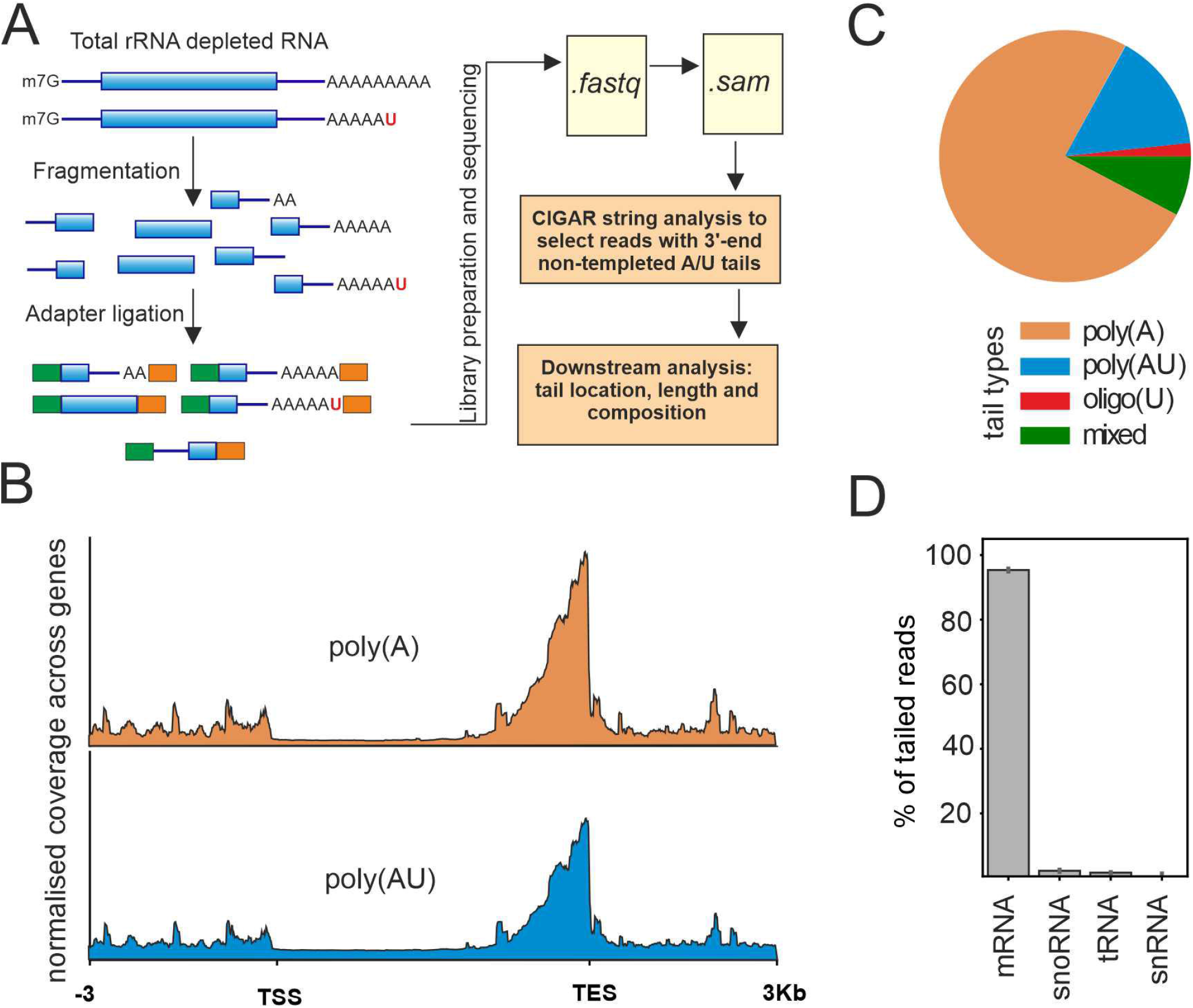
Identification of uridylation in RNA-seq data. **(A)** Scheme of workflow used to identify uridylation. Libraries prepared by ligating adapters to fragmented RNA contain 3’-end uridylated reads. We searched for such reads using information encoded in the “.sam” file CIGAR string. **(B)** The majority of detected reads with un-templated A/U extensions map closely to annotated transcription end sites (TES). **(C)** Using our approach to search for reads with A/U un-templated extensions we detected reads with pure poly(A) or poly(A) extensions additionally uridylated at the 3’-end - poly(AU). Additionally, some reads had pure U tails - oligo(U) or did not fall into any of those categories - mixed. **(D)** The majority of detected reads with un-templated A/U extensions longer than 4nt mapped to mRNAs.

As expected, the identified reads mapped to the 3’-end of RNAs (Fig. 1B). In agreement with our hypothesis in the wild-type strain, we detected reads with poly(A) and poly(AU) extensions. Those were two dominant categories of detected tailed reads, accounting for 90 % of all A/U tailed reads (Fig.1C). Both poly(A) and poly(AU) reads mapped at the 3’-end of RNA confirming that we observe predominantly uridylation of pre-existing poly(A) tails (Fig.1B). Oligo(U) reads were detected at a much lower frequency (Fig. 1C). We also recorded mixed reads where the order of As and Us did not fit any of the above-mentioned categories, however, we suspect that the majority of those originate from mismatches at the boundary site. As expected, the detected tailed reads aligned predominantly to mRNAs, and we further focused our analysis on this category (Fig. 1D).

### Both Cid1 and Cid16 participate in the uridylation of *S. pombe* mRNA

To confirm that observed uridylation originates from the enzymatic activity of terminal uridyltransferases, we created deletion strains lacking either or both *cid1* and *cid16*, encoding the known fission yeast TUT-ases. Sequencing libraries were prepared in the same way as for the wild-type cells, followed by detecting reads with the 3’-end A/U non-templated extension in the sequencing data. In agreement with uridyltransferases function, the detected mRNA uridylation decreased in TUT-ase deletion strains (Fig.2A). This result confirms that our data reflect the underlying biology and validates our method of detecting uridylation.

The *cid1* deletion resulted in the loss of 61.4% of uridylation compared to the wild-type cells (12.7% uridylated reads in wild-type in the comparison with 4.9.0% in *∆cid1*). Deletion of the *cid16* gene alone had no significant impact on the uridylation frequency. Notably, deletion of both *cid1* and *cid16* resulted in further loss of uridylation (79.5% loss compared to the wild-type cells) (Fig. 2A). Therefore, our result confirms the role of Cid1 as the main cytoplasmic uridyltransferase acting on mRNA poly(A) tails, but also show that Cid16 can participate in cytoplasmic mRNA uridylation. The Cid16 activity is only evident in the *cid1* deletion background suggesting an auxiliary role of this TUT-ase in mRNA uridylation under the condition tested.

**Figure 2.**
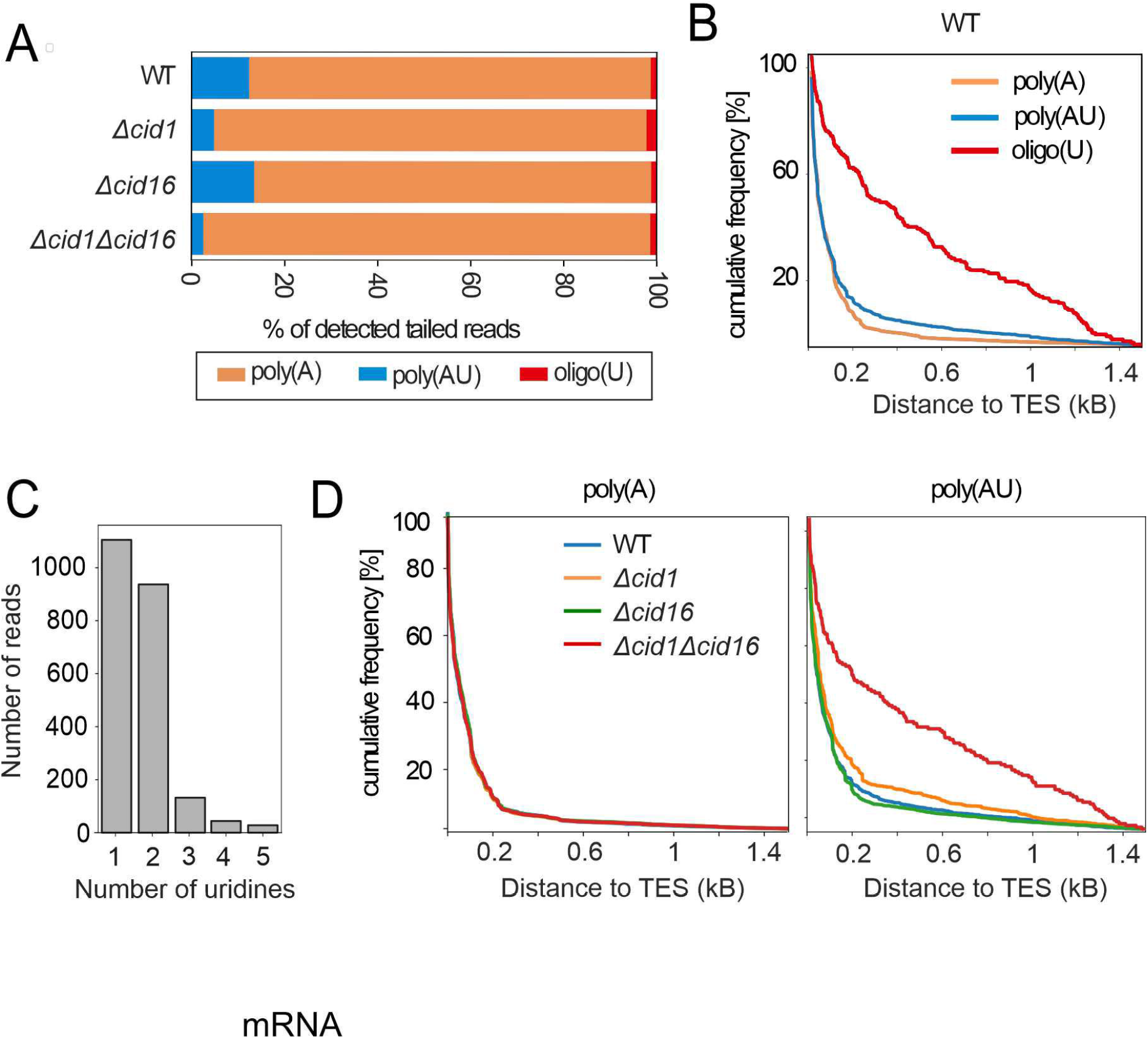
Both fission yeast uridyltransferases are implicated in mRNA uridylation. **(A)** Four categories of reads with A/U extensions that mapped to mRNA were identified in wild-type yeast (WT) and in single or double mutants of two annotated terminal uridyltransferases (TUT-ases) – *cid1* and *cid16*. While *cid1* deletion resulted in a significant drop in uridylation, the double mutant showed a further decrease in the number of detected uridylated reads. **(B)** In wild-type cells, reads with poly(A) and poly(AU) extensions map predominantly to annotated transcription end site (TES) while oligo(U) reads have a looser association with TES. Distance of reads from TES is shown as cumulative frequency. **(C)** Detected poly(AU) extensions carried predominantly one or two uridines. **(D)** Reads with poly(A) extensions mapped predominantly to TES in all tested mutants, while reads with poly(AU) extensions in strain with both TUT-ase deleted lost association with TES suggesting the artefactual origin of those reads.

According to the literature, uridylation occurs at the 3’-end of shortened poly(A) tails and consists predominantly of 1 or 2 uridine residues [4,5,7,16]. In agreement with this, most of the detected reads with poly(A) and poly(AU) tails mapped closely to annotated transcription end sites (TES), while this was not as clear for oligo(U) tails (Fig. 2B). Among uridylated reads detected in our data, the majority contained one or two uridines added to poly(A) tails (Fig. 2C). We noticed that while poly(A) tailed reads mapped to TES in all analysed mutants, the poly(AU) reads detected in the double mutant (*∆cid1∆cid16*) largely lost this correlation. This result suggests a possible artefactual origin of the detected residual reads indicating a lack of uridylation in the absence of both annotated uridyltransferases (Fig. 2D).

### Uridylation is pervasive in fission yeast

We next looked at individual transcripts for which we detected reads with non-templated nucleotide extensions. We took into account wild-type (WT) and *∆cid16* strains where uridylation was detected at the highest frequency. We considered transcripts for which at least 5 tailed poly(A) or poly(AU) reads were detected; this threshold resulted in 420 transcripts in the wild-type sample and 544 in the Cid16 deletion strain sample (Fig. 3A). For most of the transcripts, we detected both poly(A) and poly(AU) read categories. Namely, there were no transcripts with reads bearing only poly(AU) extensions and just a few transcripts with reads bearing only poly(A) extensions (14 and 25 transcripts for WT and *cid16*, respectively). The average read number for transcripts with only poly(A) tailed reads was low (7 reads per transcript), and there was no significant overlap between transcripts with only poly(A) tailed reads detected in wild-type cells and *∆cid16* (only one common transcript). All this suggests that uridylation is a common feature of all transcripts, and lack of detection is solely the result of a low read number.

**Figure 3.**
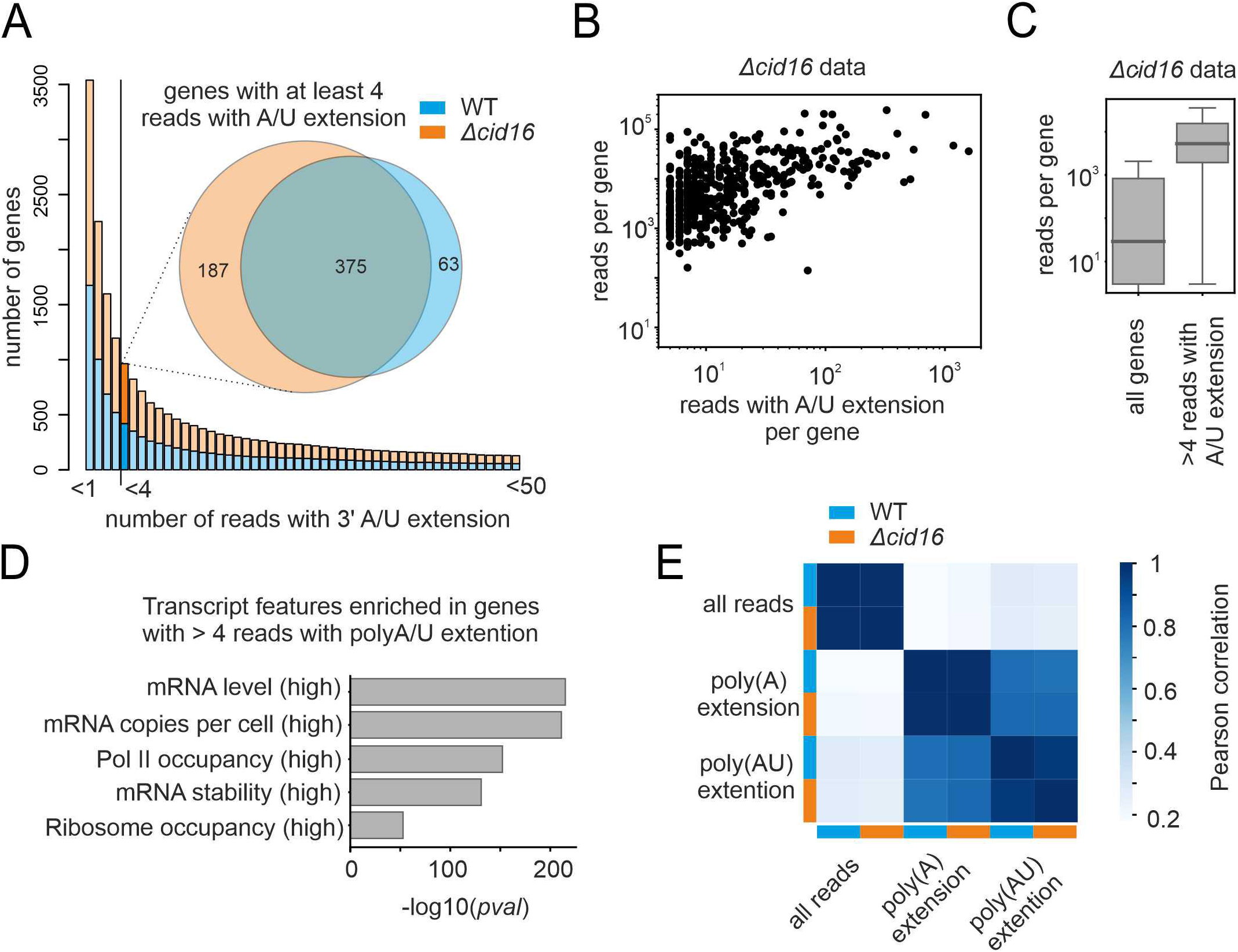
Uridylation is pervasive in fission yeast. **(A)** The number of genes with different amounts of reads with A/U extensions were plotted for two strains with high uridylation frequency (*WT* and *Δcid16*). Each bar represents the number of genes with at least as many reads detected. Venn diagram depicts overlap between genes with more than 4 A/U tailed reads detected in each strain. **(B)** The number of reads with un-templated extension positively correlates with gene expression level. Depicted are read counts for transcripts in *Δcid16* mutant plotted against the number of reads with A/U extension. Only transcripts with more than 4 reads with A/U extensions are shown. **(C)** More reads with un-templated extensions were detected for genes that are highly expressed in investigated samples. Depicted are data for *Δcid16* mutant **(D)** Transcript features of genes for which more than 4 reads with un-templated extensions were detected confirm that we observe predominantly highly expressed genes (data for *Δcid16* mutant). **(E)** Heat map of Pearson correlation calculated for different read categories detected in wild-type sample and *Δcid16* mutant – all reads, reads with poly(A) extension and reads with uridylated poly(A) extension (poly(AU)). Data for 375 genes that have more than 4 reads with A/U extensions detected in both WT and *Δcid16* samples (as shown at 3A Venn Diagram).

Accordingly, we observed that the number of reads with A/U extension positively correlated with the read number detected for a given transcript (Fig. 3B, E). Using the *cid16* deletion sample as an example (due to higher sequencing depth it had more genes passing our read number threshold), we observed that transcripts for which we detected at least 5 reads with polyA/AU extensions are the ones that are highly abundant in the sample (Fig. 3C) and are enriched in highly expressed genes categories (Fig. 3D). We thus conclude that uridylation is pervasive in fission yeast while its detection by our protocol is limited only by sequencing depth.

We observed a strong positive correlation between the number of reads with poly(A) or poly(AU) extensions detected in WT and *∆cid16* samples (Fig. 3E). There was a somewhat lower correlation between reads with poly(A) and reads with poly(AU) extensions both within and between samples (Fig. 3E). This difference might originate from different poly(A) tail lengths of different transcripts or different uridylation frequencies per transcript.

### Deletion of *cid16* but not of *cid1* leads to subtle changes in RNA steady-state levels

We observed that neither the single nor double TUT-ase deletion mutants showed significant impact on fission yeast growth in standard conditions (Fig. 4A). We next checked if phenotype of loss of uridylation can be observed when cells are confronted with unfaforable conditions that induce general stress responses [19]. All examined strains grew similarly under oxidative stress, heat shock, respiratoryconditions, in presence of respiration inhibitors, translation inhibitors and on different nitrogen sources (Supplemental_Fig_S1). We, therefore, conclude that uridylation in fission yeast is dispensable for growth control in normal conditions and is not required to stress resistance.

**Figure 4.**
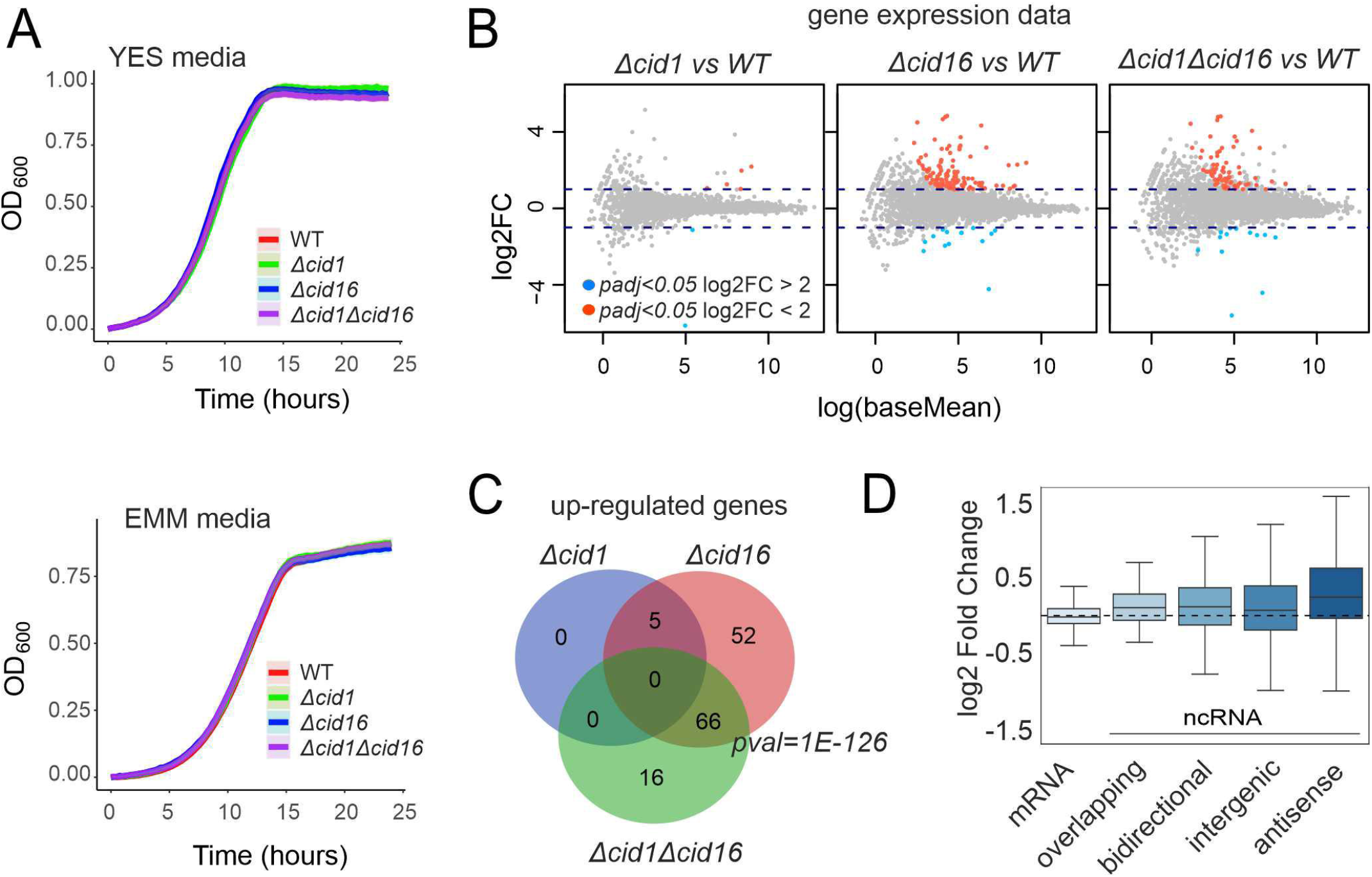
Characterisation of strains with TUT-ase deletions. **(A)** Deletion of each or both fission yeast annotated terminal uridyltransferase genes, *cid1* and *cid16*, have no detectable effect on growth. The growth of strains with deleted genes in complete (YES) and synthetic media (EMM) was recorded using Bioscreen C equipment. Growth curves represent an average of at least four repeats. **(B)** *Cid16* but not *cid1* deletion induces mild changes in gene expression. Genes expression in different TUTase deletion strains grown in YES media was compared to gene expression of the wild-type cells (WT) using DESeq2 workflow [24]. Marked red and blue on the MA plot are genes with an adjusted *p-value* below 0.05 and fold change bigger than 2 (red) or smaller than 0.5 (blue) compared to wild type. **(C)** Overlap between up-regulated genes in all three investigated strains indicate that *cid16* deletion is the main cause of observed changes. **(D)** Boxes on the plot represent changes in expression of different gene categories in *Δcid16* strain compared to wild type (WT) sample. The highest overall change was observed for antisense non-coding RNAs.

We next used our RNA sequencing data to check if eliminating uridylation results in significant changes in transcript levels. We used the standard DESeq2 pipeline to identify Differentially Expressed Genes (DEGs) in the analysed mutants compared to the wild-type strain. In agreement with the lack of growth phenotype, we observed only subtle differences between gene expression of wild-type cells and *cid1* deletion mutant, with only 7 significant DEGs between strains (*p-value* < 0.05; fold change > 2) (Fig. 4B) (Supplemental_Table_S1). We found more DEGs in *∆cid16* and double deletion strain (137 and 95 DEGs, respectively). In both strains with *cid16* deletion, we observed weak up-regulation of similar groups of genes (Fisher’s test *p-value < 2*.*2e-16*, Fig. 4C). Genes up-regulated in deletion of *cid16* or both *cid1* and *cid16* were enriched in long non-coding RNAs with 62 lncRNAs out of 132 up-regulated features in *∆cid16* strain (*p-value* = 2.4E-09), and 44 lncRNAs out of 82 up-regulated features in *∆cid1∆cid16* strain (*p-value* = 5e-08, AnGeLi) [21] (Supplemental_Table_S1). We noticed that up-regulated RNAs were predominantly antisense to other features. We used recent annotation to divide non-coding genomic features into categories: antisense, bidirectional, intergenic, and overlapping ncRNAs [22]. A comparison of overall expression changes for each category showed a general trend of antisense ncRNA upregulation in *∆cid16* (Fig. 4D). This result could be an effect of the reported function of Cid16 in uridylation of Argonaute bound small RNAs [23], which in turn could impact chromatin silencing and antisense transcription.

### Concluding remarks

We show that transcript uridylation in fission yeast shares characteristics with other species – uridylation is pervasive, it is detected almost exclusively on poly(A) tails, and the added U extensions are short. We demonstrate that, while Cid1 is the main fission yeast TUT-ase, both Cid1 and Cid16 can contribute to mRNA uridylation. Additionally, our results suggest that Cid16 can regulate non-coding antisense transcription. Our results suggest that Cid1 and Cid16 are the only uridyltransferases in fission yeast.

While uridylation of mRNAs has been reported over a decade ago, its importance for bulk mRNA decay reminds a mystery. We show here that eliminating uridylation has only a limited impact on cell growth in standard and stress conditions. On the other hand, conservation of this process suggests that it has an important role in mRNA metabolism. Fission yeast is a powerful unicellular model to study the significance of mRNA uridylation, and our data and methods lay a solid foundation for such research. Moreover, we demonstrate that transcript uridylation can be detected using simple means by analysing RNA sequencing data originating from protocols preserving the 3’-ends of RNAs. This straightforward approach will facilitate mechanistic studies of RNA uridylation also in other organisms.

## Materials and Methods

### Strains and media

*S. pombe* 972 h-strain was used as a control and to create the deletion mutants *∆cid1, ∆cid16*, and *∆cid1∆cid16. S. pombe* cells were grown in standard conditions and media (32°C in yeast extract with supplements (YES), or in Edinburgh minimal medium (EMM). To monitor respiratory growth, cells were cultivated in YE media with 2% glycerol and 0.1% glucose as carbon source. For screening responses to stressors, media were supplemented with antimycin A, chloramphenicol, CsCl, cycloheximide, ethanol, G418, H_2_O_2,_ hygromycin, KCl, MgCl_2_, NaCl, sorbitol, arginine, glutamic acid, methionine or proline (the details are provided in Supplementary Material S1.). For the heat-shock, exponentially growing cells were incubated at a temperature of 40 or 50°C for 10 or 20 minutes before re-starting growth at standard temperature.

### Growth analysis

The growth curves were obtained by monitoring changes in optical density (OD 600) using micro-bioreactor Bioscreen C. Cells were grown at 32°C in 100-well plates in 100 μL volume. Exponentially growing pre-cultures were used to start growth in the plate. The initial OD 600 of the culture was set to 0.10, and optical density was measured every 15 minutes for 48 hours.

The growth curves were extracted from the data using Pyphe growth curves Python module [4]. The maximum slope of the growth curve (describing the growth rate of the culture) and the lag phase (the time required for doubling the initial biomass) were determined. For heat map, results of the maximum slope and lag phase of deletion mutants were normalized to wild-type strain for each condition, independently. Statistical analysis was done with one-way ANOVA using Python (SciPy library). Growth curves were visualised using a custom shiny app.

### RNA-sequencing

Wild-type cells (*972 h-*) of *S. pombe* and TUT-ase deletion strains were grown in YES at 32°C to the early exponential growth phase (OD 0.5). Total RNA was isolated using the hot phenol method. RNA quality was assessed on a Bioanalyzer instrument (Agilent), treated with DNase (Turbo DNA-free, Ambion) and subsequently, 4 μg of RNA was treated with a beta version of Ribo-Zero Magnetic Gold Kit Yeast (Epicentre) to deplete rRNAs. RNA-seq libraries were prepared from rRNA-free RNA using a strand-specific library preparation protocol (a customised version of Illumina TrueSeq Small RNA) [18] and sequenced on an Illumina HiSeq instrument (126-nt paired-end reads).

Reads were checked for quality with FastQC, version 0.11.3. Adapters were removed using Cutadapt version 2.10 from Read 1 and Read 2, respectively, and then, BBMap was implemented to repair disordered reads (BBMap repair function). Reads 1 and 2 were mapped to the *S. pombe* 972 h-reference genome using HiSat2 version 2.1.0.

Count matrices were generated from BAM files using HTSeq version 0.11.1. Differentially expressed genes (DEGs) were determined using DESeq2 version 1.24.0 [24] (Supplemental_Table_S1). Based on the DESeq2 results, lists of down- and upregulated genes were prepared for each strain, and enrichments were investigated using AnGeLi (Analysis of Gene Lists) [7] with a threshold of p-value equal to 0.01 (Fisher’s exact test for count data).

### Identification of 3’ mRNA tails in RNA-seq data

The composition, length and location of 3’ non-templated extensions of sequencing reads were identified according to soft-clipping information encoded in the “.sam” file CIGAR string using a custom python script - “detect_tails_from _CIGAR.ipnyb”. It takes .sam file as an input, extracts soft-clipped reads,. Next based on CIGAR string and strand information extracts 3’-tails. Than reads are categorised as non-tailed, poly(A)-, poly(AU)-, oligo(U)-, or mixed-tailed. Finally detected tailed reads are matched to features using Bedtools v. 2.29.2 (closest function). In most of the downstream analyses, we only considered reads with extensions longer than 3 nucleotides as tailed. The exploratory data analysis was conducted with Python version 3.7.6. In the downstream analysis, we measured the distance of the last mapping nucleotide in the tailed read to annotated polyadenylation sites.

## Supplementary Data

Supplemental_Figure_S1.pdf -

Supplemental_Table_S1.xlsx - List of strains used in this study; Details of stress conditions used (corresponding to Supplementary Figure 1); List of up- and down-regulated genes detected in uridyltransferase mutants; List of significantly enriched/underrepresented transcript features categories detected among genes up-regulated in selected uridyltransferase mutants.

## Sequencing data

Raw “.fastq” files from RNA sequencing were deposited in in Sequence Read Archive (SRA, NCBI) under accession number: SUB11740974. Python script to extract tailed reads can be find in github repository - https://github.com/igib-rna-tails/. The raw data file containing reads with 3’-A/U extensions was deposited in OSF database - https://osf.io/em7vp/

## Acknowledgements

This work was funded by EU Operational Programme Innovative Economy via the Foundation for Polish Science grant FIRST TEAM awarded to M.M (POIR.04.04.00-00-4316/17). This work was supported by Wellcome Senior Investigator Award to J.B. (095598/Z/11/Z). This work was supported by National Science Center Miniatura grant (2019/03/X/NZ2/00787) awarded to L.L.-Z.. We would like to thank Szymon Świeżewski for fruitful discussions and crucial suggestions regarding data analysis.

## References

1. Parker R (2012) RNA Degradation in Saccharomyces cerevisae. Genetics 191: 671.

2. van Hoof A, Staples RR, Baker RE, Parker R (2000) Function of the Ski4p (Csl4p) and Ski7p Proteins in 3′-to-5′ Degradation of mRNA. Mol Cell Biol 20: 8230–8243.

3. Siwaszek A, Ukleja M, Dziembowski A (2014) Proteins involved in the degradation of cytoplasmic mRNA in the major eukaryotic model systems. RNA Biol 11: 1122–1136.

4. Lim J, Ha M, Chang H, Kwon SC, Simanshu DK, Patel DJ, Kim VN (2014) Uridylation by TUT4 and TUT7 marks mRNA for degradation. Cell 159: 1365–1376.

5. Rissland OS, Norbury CJ (2009) Decapping is preceded by 3′ uridylation in a novel pathway of bulk mRNA turnover. Nat Struct Mol Biol 16: 616–623.

6. Zigáčková D, Vaňáčová Š (2018) The role of 3′ end uridylation in RNA metabolism and cellular physiology. Philos Trans R Soc B Biol Sci 373:.

7. Malecki M, Viegas SC, Carneiro T, Golik P, Dressaire C, Ferreira MG, Arraiano CM (2013) The exoribonuclease Dis3L2 defines a novel eukaryotic RNA degradation pathway. EMBO J 32: 1842–1854.

8. Le Pen J, Jiang H, Di Domenico T, Kneuss E, Kosałka J, Leung C, Morgan M, Much C, Rudolph KLM, Enright AJ, et al. (2018) Terminal uridylyltransferases target RNA viruses as part of the innate immune system. Nat Struct Mol Biol 25: 778.

9. Warkocki Z, Krawczyk PS, Adamska D, Bijata K, Garcia-Perez JL, Dziembowski A (2018) Uridylation by TUT4/7 Restricts Retrotransposition of Human LINE-1s. Cell 174: 1537-1548.e29.

10. Morgan M, Kabayama Y, Much C, Ivanova I, Di Giacomo M, Auchynnikava T, Monahan JM, Vitsios DM, Vasiliauskaitŷ L, Comazzetto S, et al. (2019) A programmed wave of uridylation-primed mRNA degradation is essential for meiotic progression and mammalian spermatogenesis. Cell Res 2018 293 29: 221–232.

11. Morgan M, Much C, DiGiacomo M, Azzi C, Ivanova I, Vitsios DM, Pistolic J, Collier P, Moreira PN, Benes V, et al. (2017) mRNA 3′ uridylation and poly(A) tail length sculpt the mammalian maternal transcriptome. Nature 548: 347.

12. Thomas MP, Liu X, Whangbo J, McCrossan G, Sanborn KB, Basar E, Walch M, Lieberman J (2015) Apoptosis Triggers Specific, Rapid, and Global mRNA Decay with 3’ Uridylated Intermediates Degraded by DIS3L2. Cell Rep 11: 1079–1089.

13. Scheer H, de Almeida C, Ferrier E, Simonnot Q, Poirier L, Pflieger D, Sement FM, Koechler S, Piermaria C, Krawczyk P, et al. (2021) The TUTase URT1 connects decapping activators and prevents the accumulation of excessively deadenylated mRNAs to avoid siRNA biogenesis. Nat Commun 2021 121 12: 1–17.

14. Preston MA, Porter DF, Chen F, Buter N, Lapointe CP, Keles S, Kimble J, Wickens M (2019) Unbiased screen of RNA tailing activities reveals a poly(UG) polymerase. Nat Methods 16: 437–445.

15. Chung CZ, Jaramillo JE, Ellis MJ, Bour DYN, Seidl LE, Jo DHS, Turk MA, Mann MR, Bi Y, Haniford DB, et al. (2019) RNA surveillance by uridylation-dependent RNA decay in Schizosaccharomyces pombe. Nucleic Acids Res 47: 3045–3057.

16. Chang H, Lim J, Ha M, Kim VN (2014) TAIL-seq: Genome-wide determination of poly(A) tail length and 3’ end modifications. Mol Cell 53: 1044–1052.

17. Schlackow M, Marguerat S, Proudfoot NJ, Bähler J, Erban R, Gullerova M (2013) Genome-wide analysis of poly(A) site selection in Schizosaccharomyces pombe. RNA 19: 1617–1631.

18. Malecki M, Bitton DA, Rodríguez-López M, Rallis C, Calavia NG, Smith GC, Bähler J (2016) Functional and regulatory profiling of energy metabolism in fission yeast. Genome Biol 17:.

19. Chen D, Toone WM, Mata J, Lyne R, Burns G, Kivinen K, Brazma A, Jones N, Bähler J (2003) Global transcriptional responses of fission yeast to environmental stress. Mol Biol Cell 14: 214–229.

20. Wang S-W, Toda T, MacCallum R, Harris AL, Norbury C (2000) Cid1, a fission yeast protein required for S-M checkpoint control when DNA polymerase delta or epsilon is inactivated. Mol Cell Biol 20: 3234–3244.

21. Bitton DA, Schubert F, Dey S, Okoniewski M, Smith GC, Khadayate S, Pancaldi V, Wood V, Bähler J (2015) AnGeLi: A tool for the analysis of gene lists from fission yeast. Front Genet 6:.

22. Atkinson SR, Marguerat S, Bitton DA, Rodríguez-López M, Rallis C, Lemay JF, Cotobal C, Malecki M, Smialowski P, Mata J, et al. (2018) Long noncoding RNA repertoire and targeting by nuclear exosome, cytoplasmic exonuclease, and RNAi in fission yeast. RNA 24: 1195–1213.

23. Pisacane P, Halic M (2017) Tailing and degradation of Argonaute-bound small RNAs protect the genome from uncontrolled RNAi. Nat Commun 8:.

24. Love MI, Huber W, Anders S (2014) Moderated estimation of fold change and dispersion for RNA-seq data with DESeq2. Genome Biol 15: 1–21.

